# Ancestral sequence reconstruction using generative models

**DOI:** 10.64898/2026.01.18.700141

**Authors:** Edo Dotan, Elya Wygoda, Asaf Schers, Iris Lyubman, Yonatan Belinkov, Tal Pupko

**Author notes:** To whom correspondence should be addressed: Tal Pupko, Yonatan Belinkov.

## Abstract

Ancestral sequence reconstruction (ASR) is a foundational task in evolutionary biology, providing insights into the molecular past and guiding studies of protein function and adaptation. Conventional ASR methods rely on a multiple sequence alignment (MSA), a phylogenetic tree, and an evolutionary model. However, the underlying alignments and trees are often uncertain, and existing models typically focus on substitutions and do not explicitly account for insertion-deletion (indel) processes. Here, we introduce BetaReconstruct, a novel generative approach to ASR that harnesses recent advances in natural language processing (NLP) and hybrid transformer architectures. Our model was initially trained on large-scale simulated datasets with gold-standard ancestral sequences and subsequently on real-world protein sequences. The reconstruction requires neither MSAs nor phylogenetic trees. We demonstrate that BetaReconstruct generalizes robustly across diverse evolutionary scenarios and reconstructs ancestral sequences more accurately than maximum-likelihood-based pipelines. We additionally provide evidence that the generative-model ASR approach is also more accurate when analyzing empirical datasets. This work provides a scalable, alignment-free strategy for ASR and highlights the ability of data-driven models to capture evolutionary signals beyond the reach of traditional methods.

## Introduction

The inference of ancestral sequences plays a crucial role in evolutionary biology, enabling the reconstruction of extinct proteins, the study of functional adaptations, and a deeper understanding of molecular evolution (Brooks & Gaucher, 2007; Chang et al., 2002, 2005; Hanson-Smith et al., 2010; Liberles, 2007). Conventional methods for ancestral sequence reconstruction (ASR) typically rely on maximum likelihood estimation to infer the most probable ancestral sequence given a phylogenetic tree, a multiple sequence alignment (MSA), and a stochastic evolutionary model (Iglhaut et al., 2024; Ishikawa et al., 2019; Jowkar et al., 2023; Koshi & Goldstein, 1996; Pupko et al., 2000, 2002; Yang et al., 1995). However, these methods are highly sensitive to errors in the alignment and to the choice of evolutionary model (Aadland & Kolaczkowski, 2020; Vialle et al., 2018).

Accurate ASR is further complicated by substantial variability in evolutionary rates and patterns across genes, lineages, and functional sites. For instance, evolutionary models differ among nuclear, mitochondrial, and plastid genomes (Abascal et al., 2007), and insertion-deletion (indel) dynamics vary considerably across phylogenetic groups (Loewenthal et al., 2021; Löytynoja & Goldman, 2008; Wygoda et al., 2024). Conventional ASR methods typically assume a single evolutionary model for all sites and branches of the tree. They also assume that sites evolve independently given the tree, in order to make inference computationally feasible. However, these simplifying assumptions are biologically unrealistic (Redelings et al., 2024; Thorne, 2000) and can introduce biases and errors in ancestral sequence inference.

Deep learning and natural language processing (NLP) techniques have emerged as powerful tools for capturing complex dependencies in biological sequences (Leclercq & Droit, 2025; Ofer et al., 2021; Rannon & Burstein, 2025; Sarumi & Heider, 2024; Shu et al., 2026). NLP-based models, particularly transformer architectures, have achieved remarkable success across a variety of sequence-related tasks, such as antibiotic resistance gene prediction (Li et al., 2021), stability prediction (Gong et al., 2023), fluorescence prediction (Wang et al., 2022) and subcellular localization prediction (Elnaggar et al., 2022). These models are trained on biological sequence datasets, enabling them to capture both contextual and evolutionary information.

Here, we present BetaReconstruct, a deep-learning framework for ASR. Our approach leverages hybrid transformer-state-space models trained on both large-scale simulated evolutionary datasets and empirical sequence data. The model directly maps unaligned sequences to ancestral sequences, bypassing the need for MSAs, phylogenetic trees, or explicit evolutionary models. This novel approach provides flexibility and allows adaptation to diverse datasets and evolutionary scenarios. Our findings suggest that NLP-based models offer a promising alternative to conventional ASR methods, providing improved adaptability across varying evolutionary dynamics. This work demonstrates the potential of deep learning to transform molecular evolution studies, enabling accurate and scalable ancestral sequence inference.

### The generative algorithm for inferring ancestral sequences

We begin by outlining how NLP approaches can be leveraged to reconstruct ancestral root sequences, by framing the problem for deep learning models. Specifically, we describe how the ancestral reconstruction task is converted to a next-token prediction task, analogous to generative modeling. In this novel formulation, the model predicts each token based on preceding ones, a common task in text and code generation (Dubey et al., 2024; Gemini Team et al., 2024; OpenAI et al., 2024). To put it differently, such models learn to complete sequences based on the context of previous tokens. Tokens serve as the “atomic” units that decompose strings into smaller segments, which are then converted to numerical vectors that the model processes during its computation (Gastaldi et al., 2025; Dotan et al., 2024; Mielke et al., 2021). In this work, tokens can represent either one or multiple consecutive amino acids.

However, a set of evolutionarily related sequences (unaligned sequences) does not naturally form a “sentence”, which is the standard input for such models. Therefore, the first step is to convert the evolutionary sequence data into a single “sentence”. Among several possible strategies (Dotan et al., 2023, 2025), we adopted the “concat” approach, which concatenates the unaligned sequences into a single “sentence”, with sequences separated by a special character, the pipe (“|”). At the end of the concatenated “sentence”, we provide a special token to mark the beginning of the prediction “<RECONSTRUCT>” (Fig. 1).

**Figure 1:**
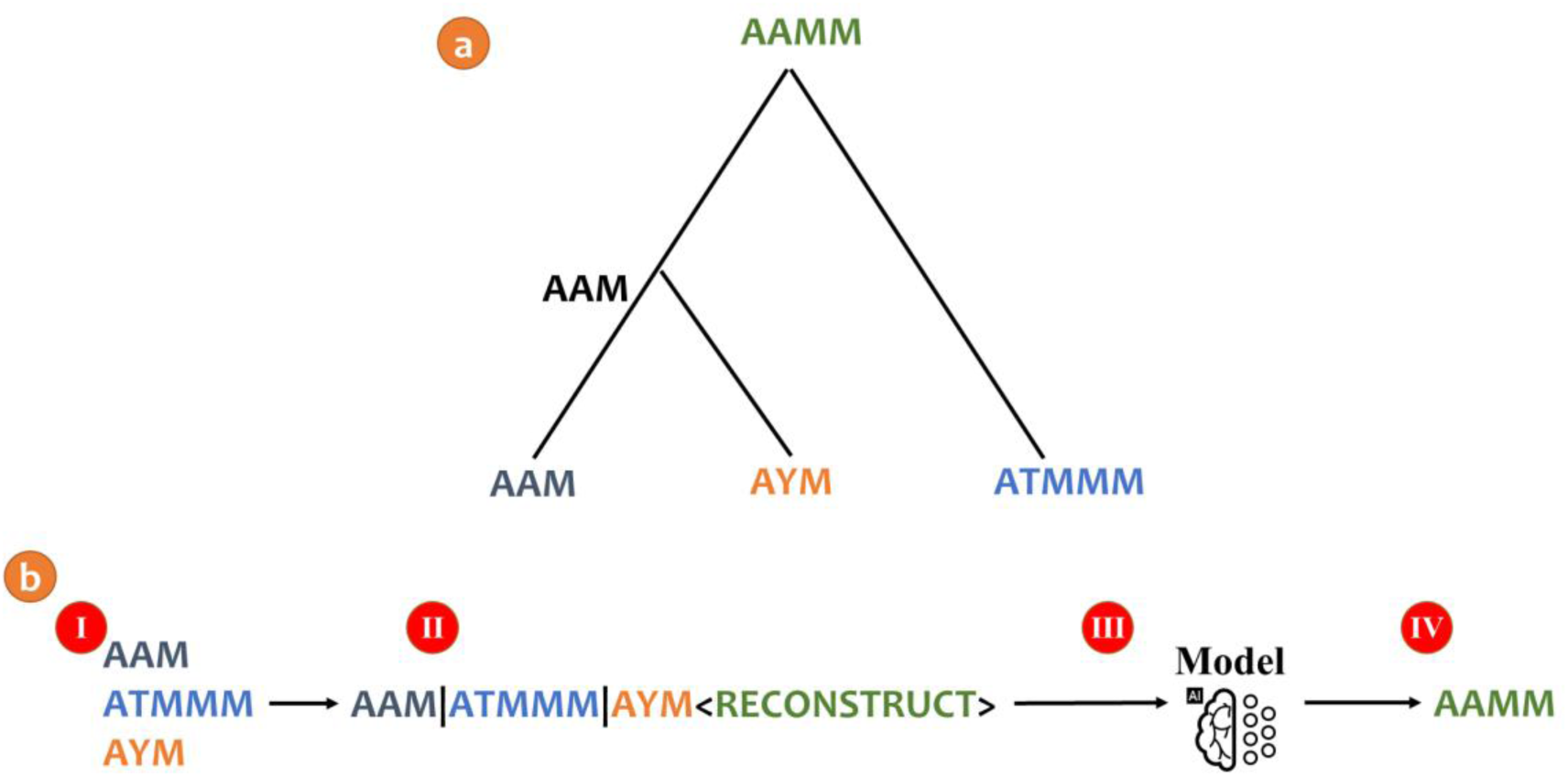
Illustration of ASR using BetaReconstruct. (a) The “true” evolutionary dynamics, in which the ancestral sequence “AAMM” evolved along a phylogenetic tree. The leaf sequences are the proteins: “AAM”, “AYM”, and “ATMMM”; (b) The BetaReconstruct pipeline: (Ⅰ) the unaligned protein sequences are provided as input; (Ⅱ) the protein sequences are concatenated with special characters between them; (Ⅲ) the model processes the input; (Ⅳ) the model generates the root ancestral sequence.

Given the special token, the model’s objective is to predict the ancestral sequence, i.e., the root sequence from which all the current sequences have evolved. Unlike traditional ASR pipelines, the input does not rely on an evolutionary model, an MSA, or a phylogenetic tree. The reconstruction is inferred directly. During training, the model is provided with unaligned sequences together with their corresponding “true” ancestral sequences, and its weights are optimized for the task (see Methods). Once trained, the model is expected to reconstruct the ancestral sequence when provided with unaligned sequences and the special token.

## Results

### Evaluation of BetaReconstruct, FastML and ARPIP

We evaluated the performance of the trained BetaReconstruct model on a simulated dataset (D7; see Supplementary Information S1), where the evolutionary model and true ancestral sequences are known. We compared the results to FastML (Pupko et al., 2000), and ARPIP (Jowkar et al., 2023), which reconstruct ancestral sequences using maximum-likelihood and also account for indel characters. We compared the following attributes: the number of matches, mismatches, and two types of gap-placement errors: false root insertions (positions erroneously included in the inferred root sequence) and false root deletions (positions erroneously deleted from the inferred root sequence). BetaReconstruct significantly outperformed FastML and ARPIP in all these attributes (paired t-test; 𝑝 < 1 × 10^−5^; Table 1). The Levenshtein distance quantifies the difference from the true ancestor, accounting for all these types of errors. The average Levenshtein distance was lower for BetaReconstruct than for FastML or ARPIP (0.18, 0.21 and 0.24, respectively; paired t-test, 𝑝 < 1 × 10^−10^). These results demonstrate the ability of BetaReconstruct to accurately predict ancestral sequences, when trained on simulated data. Notably, FastML and ARPIP require an input tree topology with its associated branch lengths for ASR. Specifically, the trees for both methods were inferred using neighbor joining. When the “true” simulated trees were provided as input for FastML or ARPIP, their performance improved dramatically, surpassing BetaReconstruct performance (see Supplementary Information S2).

**Table 1:** Comparison of BetaReconstruct, FastML, and ARPIP performance on simulated ancestral sequence data (Dataset D7; see Supplementary Information S1). The dataset includes 10–14 species, an average root sequence length of 150 amino acids, and varying insertion and deletion rates. To evaluate performance, we computed a global pairwise alignment between the inferred ancestral sequence and the true one (see Methods). Reported metrics: (1) matches, i.e., the fraction of aligned positions that match; (2) mismatches; (3) false root deletions; (4) false root insertions; (5) Levenshtein distance, i.e., the sum of mismatches and the two types of gap-placement errors; (6) Length ratio, i.e., predicted-to-true ancestor length ratio (calculated with both sequences ungapped). The different metrics in the table are statistically significant (paired t-test; 𝑝 < 1 × 10^−5^), with the exception of the length ratio metric (paired t-test; 𝑝 = 0.2).

|  | Matches | Mismatches | False root deletions | False root insertions | Levenshtein distance | Length ratio |
| --- | --- | --- | --- | --- | --- | --- |
| **BetaReconstruct** | 0.819 | 0.136 | 0.022 | 0.023 | 0.181 | 1.002 |
| **FastML** | 0.787 | 0.151 | 0.03 | 0.032 | 0.213 | 1.002 |
| **ARPIP** | 0.762 | 0.166 | 0.033 | 0.039 | 0.238 | 0.998 |

### Optimizing performance

We implemented and tested several methodological choices related to deep learning optimization, including tokenization, model architecture, training methodology, and neural network configurations. We examined tokenizers with varying vocabulary sizes and selected a tokenizer with 6,400 tokens, which provided a favorable trade-off between average token length and vocabulary size. Next, we compared the performance of three architectures: a state-space model (Mamba2; Gu & Dao, 2024), an attention-based transformer (LLAMA; Dubey et al., 2024), and a hybrid transformer (Zamba; Glorioso et al., 2024). The hybrid transformer achieved performance comparable to attention-based transformers and consistently outperformed state-space models, while requiring less memory than full-attention architectures. For each architecture, we performed a hyperparameter search to identify the configuration that maximized predictive performance. All computations were thus performed using the Zamba hybrid architecture, with a deep and narrow configuration (24 layers and a hidden vector size of 1,024). We also evaluated alternative training schemes, including multi-task learning and transfer learning, selected the best-performing model, which combined both multi-task learning and transfer learning paradigms. Detailed reports of all optimization procedures are provided in Supplementary Information S3.

### Out-of-distribution evaluation

We next evaluated two out-of-distribution scenarios, in which the test data deviated from the training distribution (Fig. 2). In the first scenario, we examined the effect of increasing the maximum branch length, i.e., the models were trained with a maximum branch length of 0.2 substitutions per site and tested on sequences with branch lengths up to 0.325 substitutions per site. We expected that performance would decline with increasing branch length, because longer branch lengths generate more sequence divergence. Indeed, the Levenshtein distance increased from 0.18 to 0.29. In addition to the increased difficulty due to higher sequence divergence, the higher Levenshtein distance can also reflect the out-of-distribution scenario. In other words, the increased error may arise from the greater difficulty of reconstructing ancestral sequences when sequences are more diverged, or because the model was not trained on highly divergent sequences. We thus tested the performance following fine-tuning, in which we trained BetaReconstruct on branches with lengths up to 0.325 substitutions per site. Our results show that the Levenshtein distance dropped to 0.28, suggesting that most of the increase in the Levenshtein distance is due to the increased difficulty and only 0.01 of the increase is due to model misspecification. In the second scenario, we assessed generalization to phylogenies containing more species than seen during training. When increasing the number of sequences from 14 to 19, BetaReconstruct exhibited only a minor increase in error (from 0.19 to 0.2). Notably, the model cannot currently handle larger numbers of sequences (tens of sequences) due to memory limitations. These results show that the models can effectively handle scenarios not encountered during training. The improved performance following fine-tuning highlights its importance for achieving better results in out-of-distribution scenarios.

**Figure 2:**
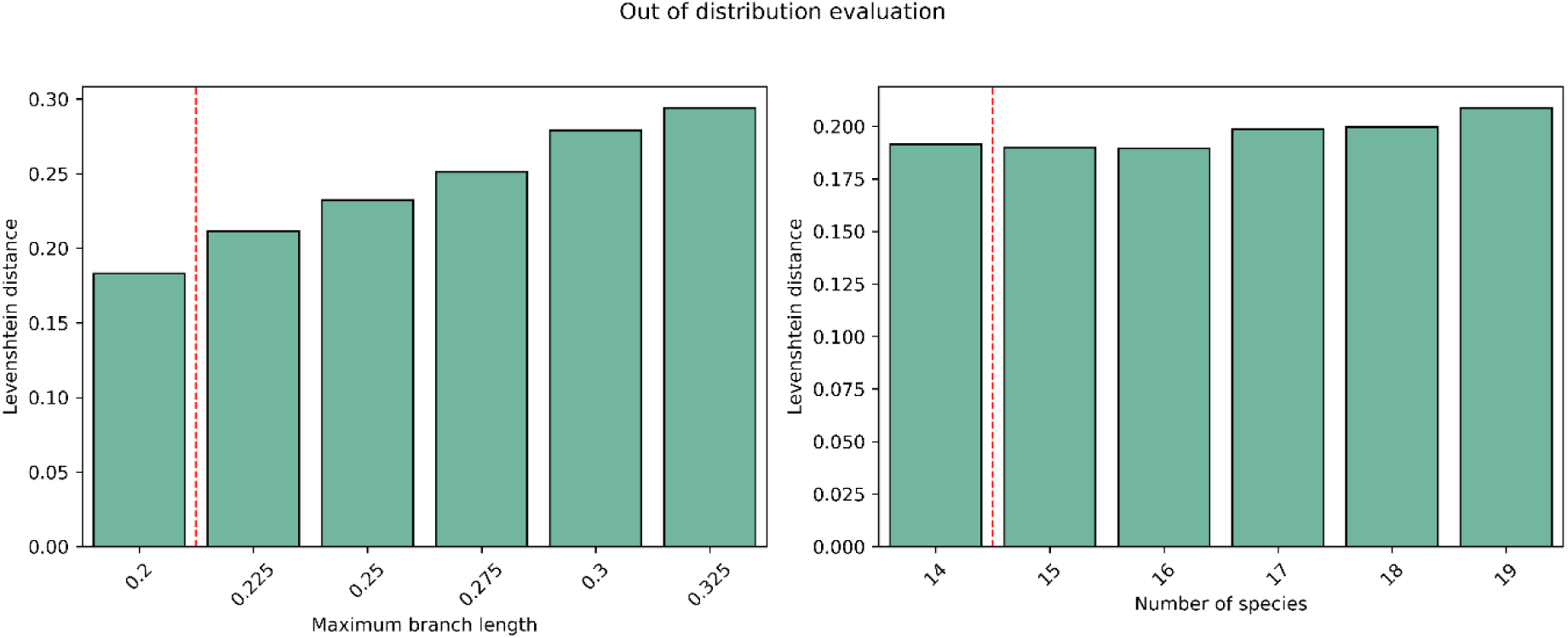
Out-of-distribution evaluation. We tested the robustness of BetaReconstruct on two types of out-of-distribution scenarios: (A) increasing maximum branch lengths beyond the training value of 0.2 substitutions per site; (B) increasing the number of sequences beyond the training value of 14. Dashed lines separate the in-distribution and out-of-distribution performance.

### Evaluating internal node prediction

For the analyses above, we provided unaligned sequences and our generative algorithm inferred the ancestral sequences at the root of the tree, without explicitly computing the underlying MSA or tree. We next studied how our model can be modified so that it can also reconstruct internal nodes, a task that is essential for elucidating the evolutionary history of specific clades for which sequence information outside the clade is available. Incorporating outgroup information further improves accuracy by providing evolutionary context and clarifying patterns of sequence divergence. Accordingly, we fine-tuned BetaReconstruct on simulated data exclusively for ancestral sequence prediction at internal nodes by explicitly marking outgroup and ingroup sequences (see Methods). This additional signal enables the model to better resolve internal ancestral states, offering a more detailed and biologically meaningful reconstruction of evolutionary paths. Table 2 compares the performance of BetaReconstruct, FastML, and ARPIP on an internal node reconstruction task using a simulated dataset, in which eight to twelve sequences form the ingroup and two sequences form the outgroup (Dataset DI; Supplementary Information S1). BetaReconstruct substantially outperformed FastML and ARPIP achieving significantly lower Levenshtein distances (0.031, 0.075, 0.104, respectively; paired t-test; 𝑝 < 1 × 10^−5^), and a higher fraction of matches (0.969, 0.925, 0.896, respectively; paired t-test; 𝑝 < 1 × 10^−5^). Notably, for FastML and ARPIP, we used the true simulated tree to extract the predicted internal node.

**Table 2:** Comparison of BetaReconstruct, FastML, and ARPIP performance in predicting internal nodes on simulated data (Dataset DI; Supplementary Information S1).

|  | Matches | Mismatches | False root deletions | False root insertions | Levenshtein distance | Length ratio |
| --- | --- | --- | --- | --- | --- | --- |
| **BetaReconstruct** | 0.969 | 0.029 | 0.001 | 0.001 | 0.031 | 1.0 |
| **FastML** | 0.925 | 0.055 | 0.01 | 0.01 | 0.075 | 1.003 |
| **ARPIP** | 0.896 | 0.051 | 0.051 | 0.002 | 0.104 | 1.062 |

### Case studies on empirical mammalian dataset

To further characterize the differences between BetaReconstruct and competing methods, we analyzed predictions for 39 mammalian protein families from a test set, i.e., these protein sequences were not included in the training and validation datasets. For each family, ancestral sequences were inferred at the root of a specific mammalian clade. We manually analyzed differences in ancestral sequences between BetaReconstruct, FastML, and ARPIP. Among these 39 predictions, 12 were identical across BetaReconstruct, FastML, and ARPIP. Out of the remaining cases, 23 predictions had an average Levenshtein distance of 0.03 between FastML and BetaReconstruct, and 0.05 between ARPIP and BetaReconstruct predictions. From the remaining divergent cases, we focused on the three most divergent ASR predictions between BetaReconstruct and FastML (protein families 5535, 353, and 7352; Levenshtein distances above 0.24), as well as the three most divergent between BetaReconstruct and ARPIP (protein families 5535, 353, and 345651; Levenshtein distances above 0.24).

Differences in indel patterns among the predicted ancestral sequences can be attributed to either erroneously inserted sequences at the root or incorrect removal of root segments. For empirical data, we do not have access to genuine ancestral sequences. However, it is possible to identify likely erroneous inferences by considering additional information from outgroup sequences, which were not used as input for the ASR algorithms.

### OrthoMaM protein family 353

Adenine phosphoribosyltransferase is a key enzyme of the purine salvage pathway that catalyzes the conversion of adenine to adenosine monophosphate, thereby contributing to nucleotide homeostasis and cellular energy balance. We reconstructed the ancestor of the *Artiodactyla* order using either BetaReconstruct, FastML, or ARPIP. As shown in Fig. 3, the different algorithms differ in their inference regarding the region spanning alignment positions 130 to 177: FastML and ARPIP suggest that this region of the MSA was absent in the ancestor of *Artiodactyla*, while BetaReconstruct indicates that it is present. However, in an outgroup clade comprising *Carnivora* sequences, this region is present, and the sequence is highly similar to that found in the *Artiodactyla* sequences. Note that the *Carnivora* sequences were not provided as input to any of these ASR algorithms. It is extremely unlikely that this region was indeed absent in the *Artiodactyla* root sequence, as this would suggest two independent insertion events of the same sequence, supporting the accuracy of BetaReconstruct relative to FastML and ARPIP.

**Figure 3:**
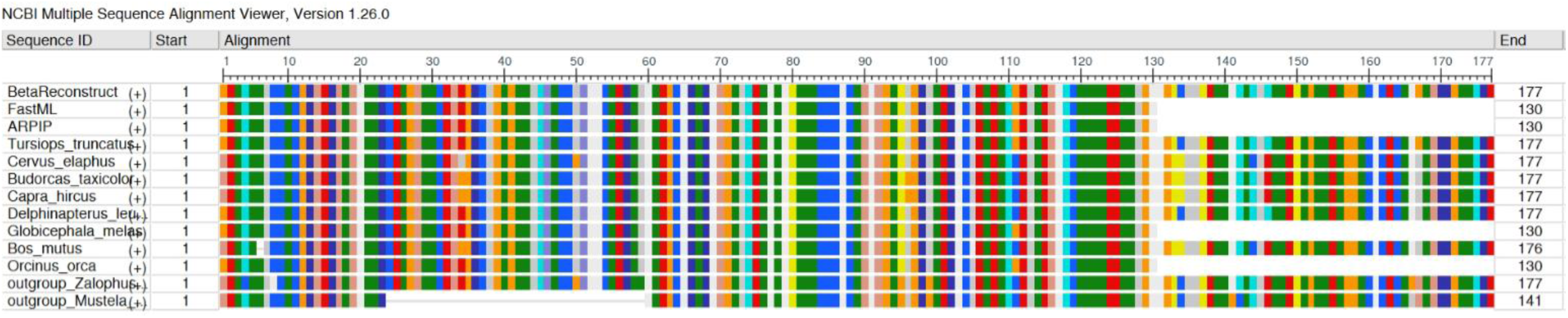
Visual alignment of the OrthoMaM protein family 353 (Adenine phosphoribosyltransferase). The first three rows are the predicted ancestral sequences of BetaReconstruct, FastML, and ARPIP. The following eight rows are the proteins used as input to the ASR pipeline, from the Artiodactyla clade. The last two rows correspond to the “outgroup” information, which are part of the Carnivora lineage. These Carnivora sequences were not used as input to the ASR pipelines. The protein sequences including the ASR predictions, the input proteins and the outgroup proteins, were aligned with MAFFT Version 7 (Katoh & Standley, 2013) and the MSA was visualized with the NCBI MSA viewer (Yachdav et al., 2016).

### OrthoMaM protein families 7352, 345651, 5535

Detailed analyses of these protein families are provided in Supplementary Information S4. Similar to protein family 353, inspection of family 7352 revealed a case in which BetaReconstruct and ARPIP inferred a deletion, whereas FastML inferred an insertion. Here, a deletion event is more consistent with the outgroup evidence. The analyses of families 345651 and 5535 were less conclusive, leaving greater ambiguity in the interpretation of the discrepancies among the ASR predictions generated by the different methods.

### Structure analysis

Finally, we predicted the ancestral root sequence of 39 protein families, each comprising about 170 sequences (the empirical test dataset). To scale ASR to datasets containing numerous species, we employed a hierarchical strategy (see Methods). We used AlphaFold 3 webserver (Abramson et al., 2024) to predict the three-dimensional protein structures of these ancestral sequences. We next computed the average structure distance between the predicted structure of the root ancestor and the predicted structure of the leaves. We expected that unrealistic reconstructions would yield predicted structures with higher structure dissimilarity, under the assumption that protein structures are generally more conserved than their sequences over evolutionary time (Illergård et al., 2009; Pascual-García et al., 2010; Zhang et al., 2010). No significant differences in average structural differences were found across the three ASR pipelines tested: BetaReconstruct, FastML, and ARPIP (see Supplementary Information S5), suggesting that these methods are comparable in terms of their ability to predict ancestral sequences that can be correctly folded and resemble the structure of leaf sequences.

### Interpreting BetaReconstruct

A notable limitation of deep learning models such as BetaReconstruct is the opacity of their decision-making process. Logit lens is an interpretability method for transformer models that allows users to “peek inside” the network at intermediate layers (Nostalgebraist, 2020). In a transformer, each layer produces an internal numerical representation (embedding) of the input sequence, but only the final layer is normally converted into predicted output symbols (in our case, amino acids). Logit lens works by taking the hidden representation at each intermediate layer and projecting it through the same output transformation used at the final layer. This makes it possible to ask what the model would predict at each layer if decoding were forced at that point (Kongmanee, 2025).

We used logit lens to analyze layer-wise probabilities, ranks, and logit values for 1,000 pairs of unaligned leaf sequences and their corresponding ancestral sequences. The results reveal a clear shift in representational behavior in the upper layers of the model (Fig. 4). The average probability assigned to the correct token remains near zero in the initial layers but rises sharply after layer 20, suggesting that the model begins to confidently consolidate ancestral sequence predictions only in the final layers. Similarly, the average rank of the correct token gradually decreases across the network, indicating a progressive refinement of sequence understanding. The logit values follow a comparable trend, increasing steadily across layers and sharply peaking toward the end, which aligns with the observed probability gain. Together, these results suggest that BetaReconstruct builds hierarchical evolutionary representations, where early layers encode broad contextual and structural information, and later layers perform the fine-grained reasoning necessary for ancestral sequence inference. These pronounced gains in rank, probability, and logit values in the upper layers may account for the superior performance of the deeper, narrower configuration (see Supplementary Information S3 for the comparison among different configurations), emphasizing the role of model depth in accurate biological sequence reconstruction.

**Figure 4:**
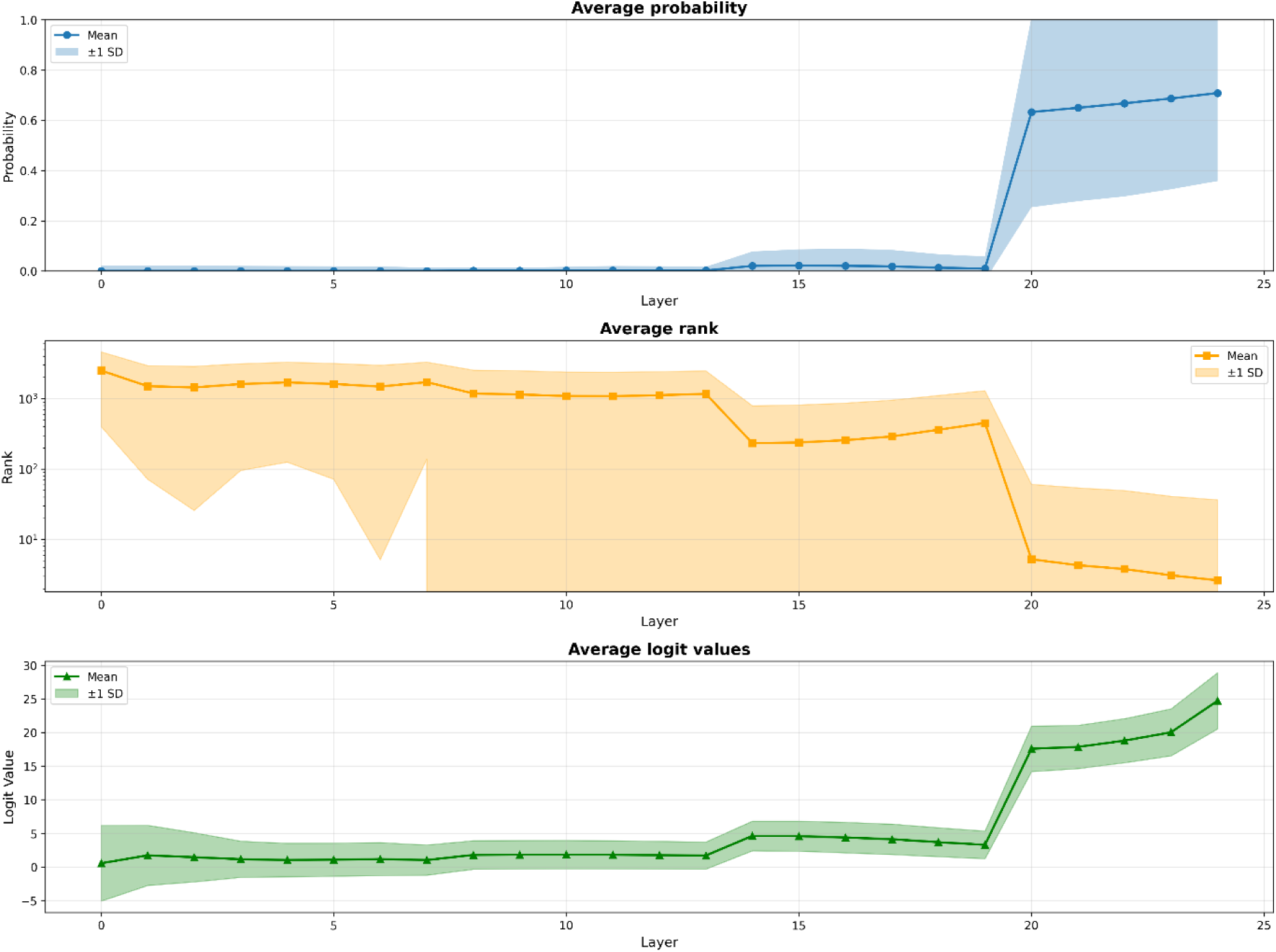
Results of the logit lens analysis applied to BetaReconstruct. The x-axis represents the transformer layer number. The three subplots display the average prediction probability (top), the average rank of the correct token (middle), and the average logit values (bottom).

## Discussion

In this work, we demonstrated how NLP techniques can be effectively applied to reconstructing ancestral sequences. By reformulating this task as a next-token prediction task, we were able to leverage advanced deep learning models to infer ancestral sequences while accounting for sequence context. Moreover, our approach bypasses traditional ASR components by directly inferring ancestral sequences from unaligned inputs, without requiring sequence alignment or phylogenetic tree reconstruction. We show that a deep-learning model trained on simulated data, and fine-tuned on mammalian sequences, performs competitively and, in some cases, outperforms established bioinformatics methods, while substantially reducing inference time. Our findings suggest a paradigm shift: rather than focusing on building specialized tools for ASR, efforts could be redirected toward developing improved simulators that accurately capture the dynamics of protein evolution. Deep learning models can then learn directly from these simulations. We hypothesize that this approach may enable accurate reconstruction even in the presence of complex evolutionary events, including large-scale rearrangements and micro-rearrangements (Pevzner & Tesler, 2003; Walker et al., 2021).

A key insight from our results is the critical role of tokenization and model architecture, especially for long biological sequences. Using a data-driven tokenizer capable of encoding multiple amino acids in a single token significantly reduced input length and, consequently, computational memory requirements. Furthermore, the hybrid architecture used for BetaReconstruct leverages the scalability and efficiency of SSMs, while preserving performance comparable to or exceeding that of attention-based models, enabling processing of much longer sequences than traditional transformers. This is particularly relevant for molecular evolution tasks such as ASR, which often involve large protein families.

Previous studies have shown that employing ensemble approaches, in which inference relies on multiple alternative MSAs rather than a single top-scoring MSA, can lead to improved inference of MSAs and tree reconstruction (Ashkenazy et al., 2019; Collingridge & Kelly, 2012; Dotan et al., 2025; Edgar, 2022). Similar ensemble-based techniques may also lead to improved ASR. In BetaReconstruct, this can be achieved by varying model configurations, permuting the order of unaligned sequences (Dotan et al., 2025), or introducing stochasticity via temperature sampling during generation (Ackley et al., 1985; Hinton et al., 2015). Two primary strategies for consolidating multiple reconstructions include majority voting (Dotan et al., 2025; Edgar, 2022) or merging (Collingridge & Kelly, 2012).

Reconstructing ancestral genomes across the tree of life presents substantial technical challenges (Garrett Vieira et al., 2020; Huang et al., 2019). Our generative approach, combined with fast and biologically faithful genome simulators, could enable ancestral genome reconstruction. To harness generative approaches for this task, several challenges remain. First, this task is complicated by the need to analyze data with a very large number of tokens (roughly proportional to the average genome length times the number of genomes). Second, for generating training and test data, realistic genome simulators are needed, with distinct evolutionary models capturing differences among genomic regions such as intergenic regions, promoters, exons, introns, enhancers, and insulators. As genomic architecture and attributes vary among clades, such a deep learning approach should be tuned to each clade, so that the method accounts for the distinct patterns of mutation, recombination, and structural variation exhibited by the different evolutionary lineages. Our leaf-training approach is particularly advantageous in this context: by learning directly on empirical genomes, the model can be tuned to account for lineage-specific evolutionary constraints, enabling more accurate and scalable ancestral genome reconstruction.

Ancestral sequence reconstruction and proto-language (ancient language) reconstruction share several conceptual and methodological parallels. Both can be formulated as a next-token prediction task, where the goal is to infer an unobserved ancestral “sequence” (protein or word) from its modern descendants. In linguistics, recent neural approaches follow exactly this paradigm: models, often transformer-based, are trained on cognate sets paired with gold protoforms, i.e., expert-validated forms (Cui et al., 2024; Kim et al., 2023; Meloni et al., 2021). Both domains rely on capturing latent evolutionary patterns. For example, the Cognate Transformer repurposes the MSA Transformer architecture, originally developed for proteins, to predict proto-language roots from cognate sets (Akavarapu & Bhattacharya, 2023). Key differences also emerge: proto-language tasks often have limited training data and shorter sequence lengths, and require careful handling of phonetic representation and alignment, while ancestral protein sequences can leverage abundant simulated data and longer alignment contexts. These parallels create a unique opportunity for shared methodological progress, where advances in one domain can seamlessly support the other.

Overall, our findings suggest that language models offer a unified, scalable, and highly generalizable framework for addressing key problems in evolutionary biology. Future extensions could explore larger models with longer context windows or multimodal architectures that integrate structural, functional, or evolutionary metadata to further boost accuracy. Incorporating explicit biological constraints, such as folding stability, structural domains, or epistatic interactions, may enable models to produce more biologically realistic ancestral reconstructions. Additionally, interpreting these models offers a unique opportunity to uncover how they internally encode evolutionary rules and to shed light on the computational strategies underlying ancestral inference.

## Supporting information

Supplementary Information: Ancestral sequence reconstruction using generative models

## Data availability

Trained models and scripts are publicly available in: https://github.com/technion-cs-nlp/BetaReconstruct.

## Materials and methods

### Outline

Our generative AI framework can be generalized to evolutionary inference tasks beyond ASR. We first describe how this framework is applied to sequence alignment and phylogenetic tree inference. Although these tasks were not the primary focus of the present study, our model was trained jointly on all three tasks: ASR, alignment, and tree reconstruction. We explain next how to formulate the three tasks as generative next-token prediction tasks. We found that this multi-task training strategy improves ASR inference accuracy. We then introduce the model architecture, a decoder-only transformer, highlighting the key design choices and optimizations for ancestral sequence generation. Next, we detail the procedures used to simulate and collect data for training and evaluation. Finally, we describe the evaluation metrics used to assess model performance.

### Generative approach for ASR

There are various deep learning models capable of generating sequences. Decoder-only models (Radford & Narasimhan, 2018) are particularly effective, producing high-quality text using next-token prediction, also known as autoregressive generation. In this work, we leverage such models to generate the ancestral root sequence. Sequence generation in a decoder-only model can be seen as a series of multiclass prediction, where the model predicts a single token at each step. The input sequence is first tokenized into substrings (represented as integers). These integers are then mapped into a high-dimensional space, termed embeddings, through an embedding layer (with dimensionality determined by the hidden state size). These embeddings are processed by a stack of decoder layers, which produce a probability distribution over the tokenizer vocabulary (via the Softmax function). A token is selected from this distribution according to a chosen generation strategy (e.g., selecting the token with the highest probability). This process repeats, until a predefined length is reached or a special end-of-sequence token (“<EOS>”) is generated.

### Multiple sequence alignment as a next-token prediction problem

To model sequence alignment, the unaligned sequences need to be converted into a single “sentence”. This is also true for the resulting MSA. The unaligned sequences are joined into a single sequence, separated by the pipe character using the “concat” format, as described for ASR. The resulting MSA is represented as a sentence using the “spaces” format, which encodes the alignment column-wise (Dotan et al., 2025). Given an MSA with *N* sequences, in the “spaces” format, the first *N* characters in the “sentence” correspond to the first column of the MSA. The next *N* characters correspond to the second column of the MSA, etc. A special token, “<ALIGN>”, is appended after the input sequence to indicate that the task is to align sequences. For the evolutionary example shown in Fig. 1a, the protein sequences “AAM,” “ATMMM,” and “AYM” are concatenated as “AAM|ATMMM|AYM<ALIGN>”. The expected output is “AAAA-Y-T-MMM-M--M-”, where each triplet corresponds to a column in the alignment: the first column is “AAA”, and the second column is “A--”, …, and the last column is “-M-”.

### Phylogenetic tree inference as a next-token prediction problem

To adapt phylogenetic tree inference to the generative model framework, we reformulated the problem as a next-token prediction task. As with the ASR and MSA tasks, the “concat” approach is used to represent the input unaligned sequences. Following the input “sentence”, we append the special token “<INFER>” to signal the beginning of the tree prediction. The tree itself is represented in a Newick format (Yu, 2022). Thus, the model learns to map between a “sentence” representing the unaligned sequences and a “sentence” (the Newick format) encoding the rooted tree topology. In the output Newick format, the leaf labels “ℓ0”, “ℓ1”, … correspond to the input sequences in order. For the example provided in Fig. 1a, the protein sequences “AAM,” “ATMMM,” and “AYM” are concatenated together with the special token “AAM|ATMMM|AYM<INFER>”. The expected output of the trained model is “((ℓ0:0.02,ℓ1:0.01):0.03,ℓ2:0.03);”, where “ℓ0”, “ℓ1”, and “ℓ2” corresponds to the sequences “AAM”, “ATMMM”, and “AYM”, respectively.

### Handling long sequences

Transformer models struggle with long sequences due to the quadratic memory requirements needed to compute attention. Consequently, these models are typically trained with a predefined maximal number of tokens, which is usually limited to a few thousand tokens (Lin et al., 2021). For the ASR task, we aim to predict the ancestral sequence of dozens of proteins, each containing hundreds to thousands of amino acids. To address this challenge efficiently, we combine two strategies: utilizing hybrid transformer architectures and tokenizers that encode multiple amino acids per token.

### Hybrid model

State space models (SSMs) have recently gained popularity due to their efficiency and scalability (Gu et al., 2022; Gu & Dao, 2024). These models leverage a compressed representation of past inputs, i.e., a state representation, whose size is linear in or independent of the input length, in contrast to the attention matrix, which scales quadratically with the number of input tokens. However, previous work has shown that attention-based transformers generally achieve higher accuracy. Hybrid models combine the strengths of both approaches by stacking SSM blocks with attention-based blocks, thereby capturing long-range dependencies while maintaining computational efficiency at scale (Glorioso et al., 2024; Lenz et al., 2024). Several hybrid architectures have been proposed, such as Zamba (Glorioso et al., 2024) and Jamba (Lenz et al., 2024). Hybrid models offer a flexible trade-off between modeling long-range dependencies and computational efficiency, making them well suited for tasks involving long biological sequences.

### Tokenization

In biological applications, deep-learning models often use tokens corresponding to single amino acids (Brandes et al., 2022; Lin et al., 2023). However, others and we have previously demonstrated that tokenizing multiple amino acids improves both performance and efficiency (Dotan et al., 2024; Lindsey et al., 2025; Sun et al., 2025). As previously mentioned, tokens are characters or substrings used to convert sequences into high-dimensional vector. These vectors are processed by the layers of the deep-learning model and are ultimately mapped back to tokens, which are then converted back to strings representing biological sequences. We trained multiple Byte-Pair Encoding (BPE) tokenizers (Sennrich et al., 2016) with vocabulary sizes ranging from 100 to 25,600 tokens. The BPE tokenizer initializes the vocabulary with the set of unique characters in the dataset, iteratively merges the most frequent token pairs, and adds the merged units as new tokens. The process terminates once the predefined vocabulary size is reached. All BPE tokenizers were trained on a subset of the training data (DT; see section ‘Data’ below). The BPE algorithm was applied to the sequence inputs of all three tasks and to the outputs of the ASR and MSA tasks. It was not applied to the output of the tree reconstruction task, which is in Newick format; this output was therefore encoded using single-character tokens. We selected a shared vocabulary size of 6,400 tokens for all tasks (see Supplementary Information S3). We note that BPE learns a data-driven vocabulary in which tokens may represent multiple amino acids or recurrent motifs, improving compression and thereby enabling the processing of longer and more numerous sequences.

### Training

We employed a two-phase training strategy, in which each phase used a distinct data type. The first phase aimed to incorporate evolutionary information by training the model jointly across the three tasks. This phase was performed using simulated data (see below). In the second phase, the model was fine-tuned specifically for ancestral sequence prediction. In both phases, model weights were optimized using a loss computed over the output tokens. Training and evaluation were implemented using the Hugging Face library (Wolf et al., 2020). During the first phase, the model was trained on “true” ancestral labels obtained from simulations. In the second phase, we introduced a novel training objective: predicting one of the descendant leaves sequences as a proxy for ancestral reconstruction when training on empirical data. Detailed training hyperparameters are provided in Supplementary Information S6.

### Transfer learning

Our training scheme extensively leverages transfer learning (Avram et al., 2024; Tan et al., 2018), allowing knowledge gained from simpler tasks to inform more complex ones. Specifically, the optimized model weights obtained from training on a simpler dataset were used to initialize the subsequent model. In phase one, we controlled dataset complexity through the simulation parameters, specifically branch length, with longer branch lengths corresponding to more challenging reconstructions. We started with a minimum branch length of 0.05 substitutions per site and increased it linearly to 0.2 substitutions per site. In phase two, the parameters optimized during phase one on simulated data were used to initialize models trained on empirical mammalian sequences.

### Computing ancestral sequences for internal nodes

In some cases, researchers are interested in reconstructing ancestral sequences at specific internal nodes of a phylogenetic tree, rather than exclusively at the root. For a given internal node, we use the term “ingroup” to refer to the leaves descended from that node and the term “outgroup” to refer to the remaining leaves. We fine-tuned BetaReconstruct (starting from the optimal weights obtained from the model trained on Dataset D7; see Supplementary Information S1) to perform ancestral sequence reconstruction at internal nodes. Specifically, we introduced special delimiter tokens to distinguish between ingroup and outgroup species: the “#” (hash) and the “|” (pipe) characters are used to enclose outgroup and ingroup sequences, respectively. For instance, a sequence enclosed by two “#” symbols is interpreted as an outgroup sequence.

### Inferring ancestral sequence for many species

To infer ancestral sequences across a large number of species, beyond what can be processed in a single model inference, we developed a hierarchical, clade-based strategy. First, we partitioned species into smaller phylogenetic clades sharing a common ancestor. We reconstructed ancestral sequences at the root of each clade, and these inferred sequences subsequently served as inputs at the next hierarchical level, allowing us to iteratively infer ancestors at progressively deeper evolutionary time points. In practice, this iterative procedure was repeated three to four times for each of the mammalian proteins analyzed. This scalable approach enables the analysis of large and complex datasets.

### Data

We used two types of data in this study: simulated protein sequences (phase one training) and empirical mammalian protein sequences (phase two training). Simulated data enabled us to directly incorporate evolutionary principles into the training process. The key advantage of using simulations is that the “true” labels are known, enabling fully supervised training and accurate evaluation. Moreover, the quantity of simulated data is effectively unlimited, allowing for extensive experimentation across a wide range of evolutionary scenarios. However, the main limitation is that the evolutionary models used for simulation are simplified representations of real molecular evolution. As a result, they do not fully capture complex factors arising from selective constraints imposed by protein structure and function, such as dependencies among alignment positions and variation in evolutionary processes across sites and tree branches. To address this limitation, we supplemented training with empirical protein sequence data from diverse mammalian species. Since ground-truth ancestral sequences are unavailable, the model was trained to predict one species from the others without relying on a multiple sequence alignment or an explicit phylogenetic tree. This dual training strategy, grounded in simulations but informed by real data, enables the model to leverage both theoretical evolutionary knowledge and empirical biological diversity.

### Simulated data

To train and evaluate BetaReconstruct across ASR, MSA, and phylogenetic tree inference tasks, we generated large synthetic datasets consisting of unaligned sequences together with their corresponding ground-truth alignments, ancestral sequences, and phylogenetic trees (Wygoda et al., 2025). In each dataset, sequences evolved along randomly generated phylogenetic trees with diverse topologies and branch lengths. The simulator allows insertions and deletions to evolve with independent rates and length distributions, which is critical for realistically capturing molecular evolutionary complexity (Loewenthal et al., 2021). From each simulation, we extracted a pair consisting of a set of extant sequences and one of the following: the ancestral sequence at the root, the MSA, or the phylogenetic tree. These ground-truth labels enabled consistent supervised training and rigorous evaluation of model performance. Detailed simulation parameters are provided in Supplementary Information S1.

### Mammalian data

To train and evaluate BetaReconstruct on real biological data, we used OrthoMaM version 12, a curated dataset of orthologous genes from a broad range of mammalian species (Allio et al., 2024). We extracted multiple unaligned protein sequences across diverse mammals and grouped them into clades based on taxonomic orders: *Rodentia*, *Primates*, *Chiroptera*, and *Carnivora*. Each clade was used to fine-tune BetaReconstruct on one representative species (leaf) randomly selected from the group. The training set comprised 119,064 unaligned proteins, each paired with a randomly selected leaf. The OrthoMaM database contains approximately 15,000 proteins, and multiple training examples can be generated from a single protein and order, depending on the number of species represented per protein. The validation set includes 16,240 data points of the *Artiodactyla* order. Finally, we evaluated the performance on a test dataset of 39 OrthoMaM protein families that were entirely excluded from the training data (see Supplementary Information S1).

### Evaluating performance

We monitored the cross-entropy loss (Szegedy et al., 2015) on both the training and validation datasets. For each token in a sequence, the model outputs a probability distribution over the full tokenizer vocabulary. The cross-entropy loss quantifies how well this predicted distribution aligns with the actual (true) distribution, where the actual token probability is one and all other tokens are zero. Formally, it is computed as the negative logarithm of the predicted probability assigned to the correct token. The sequence loss is obtained by averaging the value across all predicted tokens in the sequence. The dataset loss is computed by averaging the loss across all sequences. Lower loss indicates that the model is assigning higher probabilities to the correct tokens and therefore making more accurate predictions. The training loss is computed on the training data and used to update the weights of the model. The validation loss is computed on the validation data and used to select hyperparameters, track performance and avoid model overfitting.

To directly assess the accuracy of inferred ancestral sequences, we compared them against the “true” ancestral sequences (when the data are simulated). The predicted sequence was globally aligned to its corresponding true sequence using a simple pairwise alignment scheme with a scoring system of +1 for matches and –1 for mismatches or indels. From these alignments, we extracted several evaluation metrics: (i) the number of exact matches; (ii) the number of characters present in the true ancestral root sequence and missing from the inferred root sequence, i.e., false root deletions; (iii) the number of characters absent in the true ancestral root sequence and present in the inferred root sequence, i.e., false root insertions; and (iv) the number of mismatches between the two sequences. In addition, we computed the Levenshtein distance (Wagner & Fischer, 1974), defined as the minimum number of substitutions, insertions, or deletions required to transform a sequence into another one, and equal to the sum of metrics ii, iii, and iv. Thus, the Levenshtein distance provides a global summary of reconstruction errors, integrating all types of errors into a single score. Together, these measures enable a detailed characterization of the accuracy of the inferred ancestral sequence across models and configurations. We normalized the metrics by dividing the score by the length of the true ancestral sequence. If a metric score was higher than one, we set it to one, and thus the distance is a number between zero and one.

### Logit lens

The logit lens is a mechanistic interpretability method used primarily for analyzing transformer-based models. It projects the intermediate hidden representations (i.e., embeddings), from each layer directly into the final output vocabulary space using the vocabulary matrix (Nostalgebraist, 2020). By doing so, we can effectively “peek” at the model’s partial predictions at each processing stage (Kongmanee, 2025). Logit lens traces how predictions evolve through the network layers, often revealing a clear trajectory from initial broad or even nonsensical guesses to a refined and confident final prediction in the later layers. This allows researchers to localize where specific computations or reasoning steps are occurring within the network architecture. All logit-lens computations were done using the Hugging Face library (Wolf et al., 2020).

