## Supplementary Information: Ancestral sequence reconstruction using generative models for "Ancestral sequence reconstruction using generative models"

[1]The Shmunis School of Biomedicine and Cancer Research, George S. Wise Faculty of Life Sciences, Tel Aviv University, Tel Aviv 69978, Israel.

[†]To whom correspondence should be addressed:

### Supplementary Information

#### Supplementary Information S1: Specific datasets and simulation parameters

Sequences were simulated along a tree according to the following procedures. We first sampled a random phylogenetic tree with 10 to 14 species, using the ETE3 library with default parameters (Huerta-Cepas et al., 2016). We sampled the indel parameters, based on the following ranges: $R_i, R_d \in (0.0, 0.05)$, and $A_i, A_d \in (1.01, 2.0)$, where $R_i, R_d, A_i, A_d$ corresponds to the insertion rate, deletion rate, a parameter for the insertion Zipfian distribution, and a parameter for the deletion Zipfian distribution, respectively (Loewenthal et al., 2021). Insertions and deletions were sampled independently, allowing for a rich-indel model. Regarding the substitution replacement model, we used the "WAG+G" model with the gamma alpha parameter equal to 1.0 and four discrete gamma categories. Each such simulation instance provided the following data: (a) true tree, including topology and branch lengths; (b) a set of unaligned sequences, including unaligned ancestral sequences; (c) the true alignment.

We first generated D1, consisting of 1,080,000 simulation instances, which were split into 1,050,000 instances for training and 30,000 instances for validation. The branch lengths in these cases were relatively short (between 0 and 0.05 substitutions per site), making the inference of alignments and ancestral sequences easy (Table S2). When training on D1, we established pairs (*X*, *Y*), where *X* is the set of unaligned sequences, and *Y* is the label. For one-third of the simulation instances in each dataset, the *Y* label was the unaligned root ancestral sequence. For another one-third, the *Y* label was the multiple sequence alignment, and for the remaining instances, the *Y* label was the string representing the tree in Newick format.

We next generated D2 by generating 1,080,000 additional simulation instances, this time with longer branches (Table S2). The optimized weights of the model that was trained on D1 were the starting point for the model trained on D2. This process was repeated until the model was trained on D1, D2, …, D7, which increase in difficulty (Table S2).

We generated an additional dataset, DT ("dataset tokenizer"), which was used to train and evaluate the tokenizer. The main difference between DT and the other datasets is the root sequence length, which was sampled from a normal distribution with a mean of 345 and a standard deviation of 76. These distribution parameters were derived from a large dataset of empirical proteins (Nevers et al., 2023). For the DT, we generated 7,000 simulation instances, which were split into 6,000 training instances and 1,000 validation instances.

While datasets D1-D7 were used for the task of predicting the root sequence, we also generated Dataset DI ("dataset internal nodes"), on which we trained the model to predict not only the root sequence but also the ancestral sequence at internal nodes. In DI, eight to twelve species were used for the ingroup sequences and two for the outgroup sequences. Additionally, we simulated another dataset for out-of-distribution evaluation. The out-of-distribution dataset includes only 5,000 validation instances (as no training was performed), with branch lengths gradually increasing, starting at 0.225 substitution per site and increasing it up to 0.325. Similarly, we simulated 5,000 validation instances with an increasing number of unaligned sequences, starting at 15 sequences and increasing up to 19.

For the mammalian empirical analysis, we used the OrthoMaM database (version 12; Allio et al., 2024). The test dataset consisted of 39 protein families, specifically: 100188893, 10815, 121214, 1487, 2170, 2331, 29923, 353, 51077, 64429, 10468, 11345, 121355, 1984, 22954, 283297, 345651, 388394, 55929, 8666, 6222, 5213, 55379, 5236, 6472, 55207, 5535, 4975, 151242, 1445, 7389, 7476, 84639, 977, 7352, 8078, 7345, 9978, and 84649.

*Table S2: Specific parameters for the simulated datasets. Parentheses indicate ranges. Root lengths were sampled from a normal distribution with mean μ and standard deviation σ.*

| Dataset | Branch lengths | Number of species | Root length |
|---|---|---|---|
| **D1** | (0, 0.05) | [10, 14] | $\mu = 150, \sigma = 30$ |
| **D2** | (0, 0.075) | [10, 14] | $\mu = 150, \sigma = 30$ |
| **D3** | (0, 0.10) | [10, 14] | $\mu = 150, \sigma = 30$ |
| **D4** | (0, 0.125) | [10, 14] | $\mu = 150, \sigma = 30$ |
| **D5** | (0, 0.15) | [10, 14] | $\mu = 150, \sigma = 30$ |
| **D6** | (0, 0.175) | [10, 14] | $\mu = 150, \sigma = 30$ |
| **D7** | (0, 0.20) | [10, 14] | $\mu = 150, \sigma = 30$ |
| **DT** | (0, 0.20) | 14 | $\mu = 345, \sigma = 76$ |
| **DI** | ingroup: (0, 0.08), outgroup: (0.1, 0.35) | ingroup: [8, 12], outgroup: 2 | $\mu = 150, \sigma = 30$ |

#### Supplementary Information S2: ASR Performance on simulated data in cases where the true phylogenetic tree is given as input to FastML and ARPIP

We evaluated the performance of FastML and ARPIP when given the "true" tree as input. The results for BetaReconstruct reported in Table S1 could not easily incorporate this information and would require additional specialized training. An example of such specialized training (where data are simulated using a specific tree with outgroup of two species) is provided by training BetaReconstruct to reconstruct internal nodes rather than the root node. In contrast, both FastML and ARPIP showed a dramatic improvement.

*Table S1: ASR performance when the "true" phylogenetic tree is provided as input for FastML and ARPIP (Dataset D7).*

| | Matches | Mismatches | False root deletions | False root insertions | Levenshtein distance | Length ratio |
|---|---|---|---|---|---|---|
| **BetaReconstruct** | 0.819 | 0.136 | 0.022 | 0.023 | 0.181 | 1.002 |

| | | | | | | |
|---|---|---|---|---|---|---|
| **FastML** | 0.891 | 0.085 | 0.014 | 0.01 | 0.109 | 1.007 |
| **ARPIP** | 0.887 | 0.087 | 0.016 | 0.011 | 0.113 | 1.006 |

### Supplementary Information S3: Optimization

#### Effect of tokenizer size on efficiency

In the context of protein sequence analysis, tokenization transforms the protein sequences into integers, called tokens, where a token can represent a single amino acid or a short k-mer. Notably, when tokenizing a single protein sequence, tokens may represent one or more consecutive amino acid positions. The BPE (Sennrich et al., 2016) tokenizer takes as input a set of sequences and determines the optimal way to define tokens, given a constraint on the total number of tokens, known as the vocabulary size. For a specific set of unaligned sequences, the vocabulary size determines the number of tokens present in the set (roughly equivalent to the number of words in a given text). When the vocabulary size is high, the average token captures a longer stretch of amino acids, which reduces the number of tokens that need to be processed by the model. The resulting fewer tokens reduces the memory footprint. However, selecting a vocabulary size too large may reduce performance, because the prediction model would have to select next tokens from a wider distribution. We tested the BPE tokenizer with various vocabulary sizes: 100, 200, 400, 800, 1,600, 3,200, 6,400, 12,800, and 25,600 on simulated data (Dataset DT; see Supplementary Information S1). Fig. S1 reports the average number of tokens across the unaligned sequences in this dataset. This average number of tokens plateaus as the vocabulary size increases beyond 6,400 tokens. Based on this observation, we chose a vocabulary size of 6,400 for all models, resulting in a token representing on average 3.82 amino-acids.

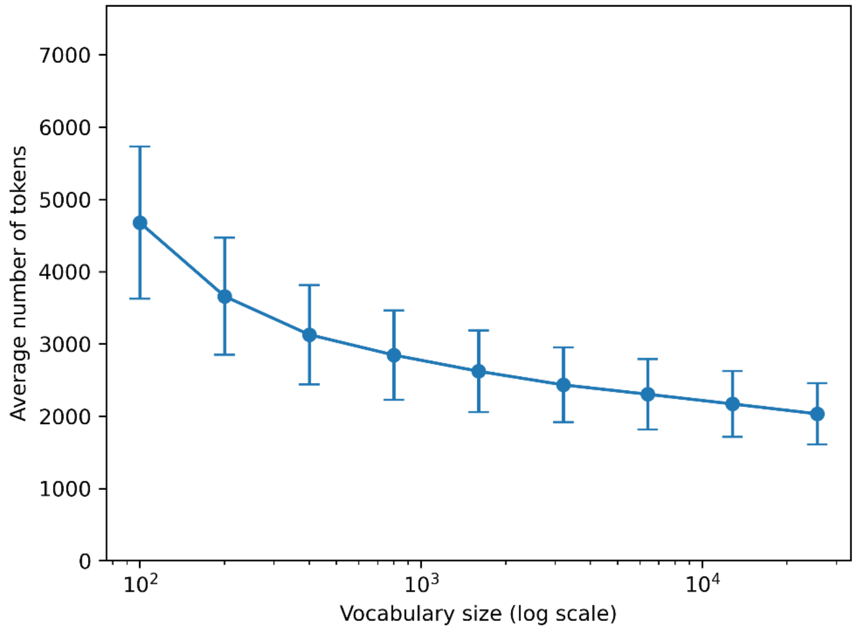


*Figure S1: Effect of tokenizer vocabulary size on the average number of tokens in Dataset DT.*

#### Effect of architecture on performance

We tested three different transformer architectures: Zamba (Glorioso et al., 2024), LLAMA (Dubey et al., 2024), and Mamba2 (Dao & Gu, 2024). Mamba2 represents a state-space model, LLAMA is a purely attention-based model, and Zamba is a hybrid architecture combining state-space and attention layers (see Methods). Hyperparameters were optimized for each architecture (see below). Performance comparisons revealed notable differences across architectures (Table S3). Zamba and LLAMA consistently achieved better performance than Mamba2, suggesting that attention layers contribute to improved performance. The hybrid model performed similarly to the attention-based model in terms of Levenshtein distance (0.06 vs. 0.058), while maintaining lower memory usage and faster training times. Consequently, the Zamba architecture was used in all subsequent analyses.

*Table S3: Effect of architecture on validation loss and ASR performance. We tested three architectures: hybrid (Zamba), attention-based (LLAMA), and state-space (Mamba2). All models contain approximately 200 million parameters.*

| **Architecture** | Zamba | LLAMA | Mamba2 |
|---|---|---|---|
| **Validation loss** | 1.248 | 1.405 | 6.656 |
| **Average Levenshtein distance** | 0.06 | 0.058 | 0.91 |

#### Hyperparameter tuning for each architecture

The above comparison among architectures necessitates finding optimal hyperparameters for each architecture. Thus, we considered three network configurations, and two learning rates per architecture type (Table S4). Since this grid-search training is extensive, we trained the models on D1 for 5,120 steps. The Mamba2 models exhibited substantially higher validation losses and error rates across all configurations, suggesting instability or incompatibility of this architecture under the current training setup, and hence we did not consider this architecture further. Across all three configurations, when selecting the learning rate that led to the highest performance, Zamba outperformed LLAMA in terms of validation loss (1.367 vs. 1.535, 1.248 vs. 1.405, 1.419 vs. 1.436 for Configurations 1, 2, and 3, respectively). In terms of Levenshtein distance, LLAMA had slightly better performance than Zamba for Configurations 2 and 3, but substantially less so in Configuration 1. Due to the higher memory requirements and longer training times compared to Zamba, we adopted the Zamba architecture for BetaReconstruct.

*Table S4: Evaluation of architecture, configuration, and learning rate combinations. Models were compared using two metrics: validation loss and average Levenshtein distance. All 18 architecture–configuration–learning-rate combinations contained approximately 200 million parameters. Each model was trained on Dataset D1 for 5,120 steps.*

| | Configuration 1 (deep and narrow) | | | Configuration 2 (balanced) | | | Configuration 3 (shallow and wide) | | |
|---|---|---|---|---|---|---|---|---|---|
| **Architecture** | Zamba | LLAMA | Mamba2 | Zamba | LLAMA | Mamba2 | Zamba | LLAMA | Mamba2 |
| **Layers** | 24 | 24 | 24 | 16 | 16 | 16 | 12 | 12 | 12 |
| **Hidden size** | 1,024 | 832 | 1,024 | 1,280 | 1,024 | 1,280 | 1,408 | 1,152 | 1,536 |
| **Validation loss (learning rate of $1 \times 10^{-4}$)** | 1.367 | 1.535 | 8.296 | 1.248 | 1.405 | 7.661 | 1.419 | 1.436 | 7.896 |
| **Average Levenshtein distance (learning rate of $1 \times 10^{-4}$)** | 0.065 | 0.998 | 1.0 | 0.06 | 0.059 | 1.0 | 0.065 | 0.059 | 0.929 |
| **Validation loss (learning rate $1 \times 10^{-5}$)** | 2.383 | 5.719 | 6.793 | 2.237 | 5.631 | 6.656 | 2.417 | 5.567 | 7.313 |
| **Average Levenshtein distance (learning rate $1 \times 10^{-5}$)** | 0.115 | 1.000 | 0.911 | 0.101 | 0.802 | 0.99 | 0.075 | 0.800 | 0.959 |

#### Comparing Zamba Configurations

We continued training the three Zamba configurations on datasets with progressively increasing complexity. The results presented for the most complex dataset (D7) show clear differences in model performance. As shown in Fig. S2, Configuration 1 outperformed the others across both validation loss and Levenshtein distance. These results indicate that deeper and narrower architectures generally yield more accurate predictions. Notably, although the difference in validation loss between Configurations 1 and 3 was relatively small (0.5%), their ASR performance diverged substantially (8.5%). This emphasizes the importance of assessing models using biologically meaningful metrics rather than relying solely on validation loss.

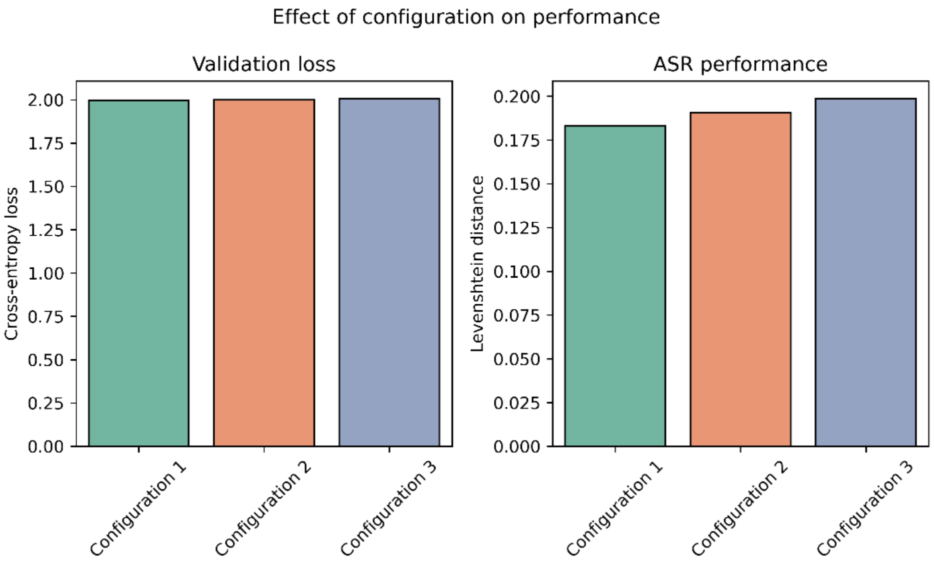


*Figure S2: Evaluating different configurations on a simulated dataset (D7). Configurations 1, 2, and 3 correspond to Zamba models with 24 layers and a hidden size of 1,024, 16 layers and a hidden size of 1,280, and 12 layers and a hidden size of 1,408, respectively. All models share a similar number of parameters, approximately 200 million.*

#### Effect of training on multiple tasks and transfer learning

To evaluate the impact of training strategies and transfer learning on model performance, we assessed three different approaches using the best-performing configuration (Configuration 1; 24 layers and a hidden size of 1,024). The first strategy, "multiple training", involved progressively training a single model across all three tasks (ASR, phylogenetic tree inference, and MSA) starting with simpler datasets and gradually increasing complexity (see Methods). The second approach, "specific training", trained a separate model for the ASR task only, using datasets with gradually increasing complexity. The third approach, "without transfer learning", involved training a model from scratch on the most complex dataset. As shown in Fig. S3, the model that leveraged transfer learning outperformed the model trained without it (error rate of 0.18 vs. 0.22). Although "specific training" provided moderate performance, it remained inferior to "multiple training", highlighting the advantages of joint training and knowledge transfer for learning generalizable representations of evolutionary patterns.

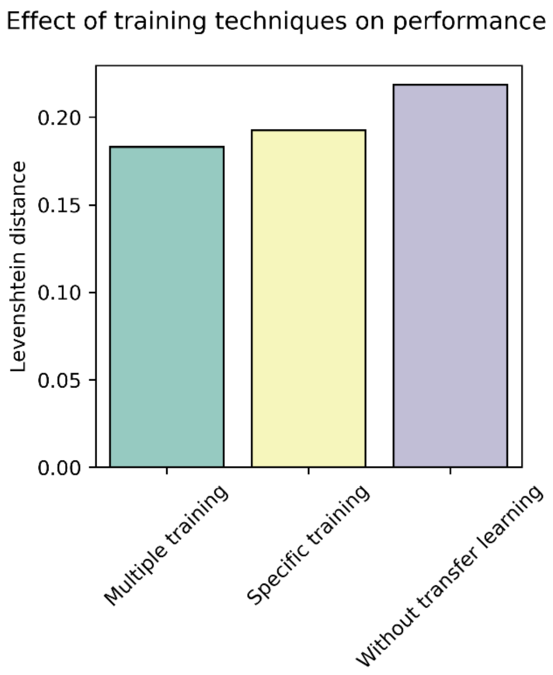


*Figure S3: Effect of training strategy on model performance. We compared three training strategies across the ASR inference task. "Multiple training" refers to a single model trained progressively from simple to complex datasets across the three tasks (ASR, MSA, and phylogenetic inference). "Specific training" involves gradually training a separate model for the ASR task only, with datasets of gradually increasing complexity. "Without transfer learning" denotes a model trained from scratch (random initialization) on the full, complex dataset without exposure to simpler data. All experiments were conducted using Configuration 1.*

#### Supplementary Information S4: Detailed use cases

Here we provide detailed use cases of the top divergent predictions across our 39 test protein families. For each use case, we provide predictions from BetaReconstruct, FastML, and ARPIP (top three rows) as well as the species used to infer the ancestral sequences (from the *Artiodactyla* order). As discussed in the main paper, we also used outgroup information, i.e., proteins of a different clade (bottom two rows). We used the MAFFT alignment software (version 7; Katoh & Standley, 2013), and the NCBI MSA viewer (version 1.26; Yachdav et al., 2016) to visualize the results.

##### OrthoMaM protein family 7352

Mitochondrial transporter UCP3 is an inner mitochondrial membrane protein implicated in the regulation of proton leak, energy metabolism, and fatty acid oxidation, particularly in metabolically active tissues. Outgroup data were obtained from the order *Chiroptera*. As shown in Fig. S4, this analysis provides insights consistent with those discussed for case 353, indicating that the region spanning positions 1 to 38 was likely part of the ancestral protein of the *Artiodactyla* clade. In this case, both BetaReconstruct and ARPIP predicted the presence of this region in the reconstructed ancestral sequence.

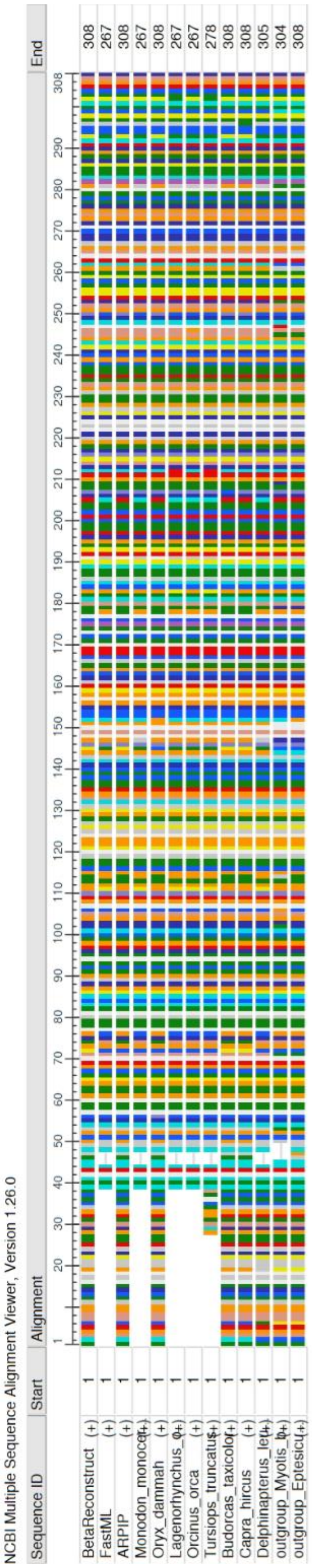


*Figure S4: Visual alignment of the protein family 7352. The first three rows correspond to the predictions of BetaReconstruct, FastML, and ARPIP, respectively. The following eight rows show the input proteins (from the Artiodactyla clade). The last two rows show the outgroup sequences, which belong to the Carnivora lineage.*

#### OrthoMaM protein family 345651

Beta-actin-like protein is a highly conserved cytoskeletal protein involved in maintaining cell structure, enabling cell motility, and supporting diverse processes such as intracellular transport and gene regulation. Unlike cases 353 and 7352, the results for this protein do not reveal a clearly inserted or deleted region, making the conclusions less definitive. A short region spanning positions 185 to 188 was present in the outgroup but absent in the ARPIP prediction (Fig. S5). Moreover, the predictions generated by BetaReconstruct and FastML were substantially more consistent with the outgroup sequences, exhibiting an average Levenshtein distance of approximately 0.04, compared with over 0.25 for ARPIP.

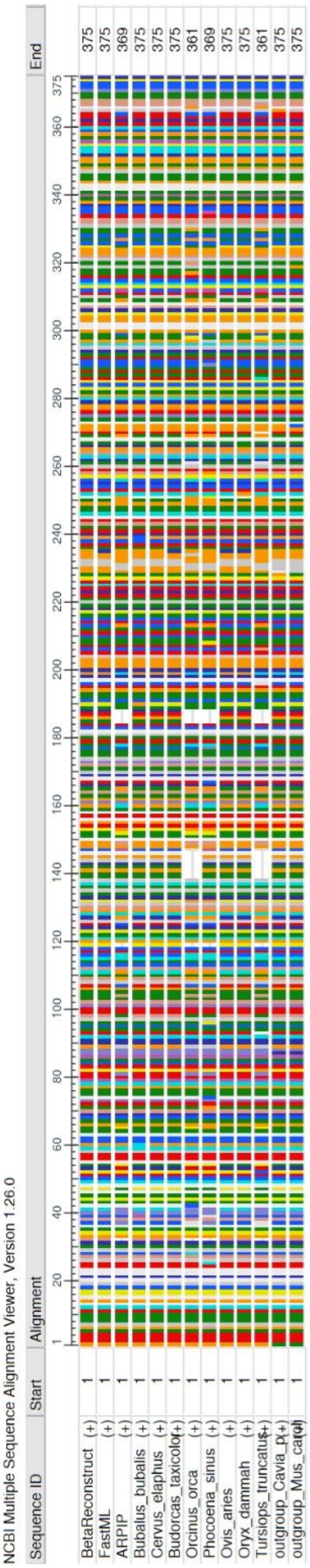


*Figure S5: Visual alignment of the protein family 345651. The last two rows correspond to the outgroup sequences, which are part of the Rodentia lineage.*

#### OrthoMaM protein family 5535

Calcineurin subunit B type 2 is a calcium-binding regulatory subunit of the serine/threonine phosphatase calcineurin, mediating calcium-dependent signal transduction pathways that regulate diverse cellular processes, including transcription and immune responses. In this example, the results are even less conclusive, as all predictions show high divergence from the outgroup sequences, with an average Levenshtein distance exceeding 0.3 (Fig. S6). In this case, ARPIP and FastML predicted identical ancestral sequences.

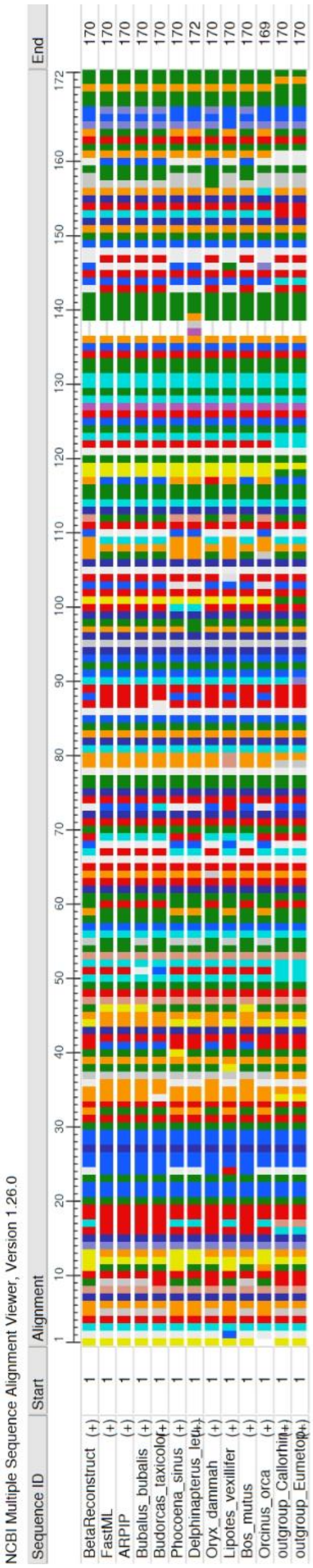


*Figure S6: Visual alignment of the protein family 5535. The last two rows correspond to the outgroup sequences, which are part of the Carnivora lineage.*

#### Supplementary Information S5: Comparing to competitors on large protein families

As no true ancestral sequences are available for real protein sequences, we evaluated the quality of the reconstructed ancestral sequences by analyzing their predicted structures. This approach is based on the observation that protein structures tend to be more conserved than their sequences during evolution (Illergård et al., 2009; Pascual-García et al., 2010; Z. Zhang et al., 2010). Consequently, an accurate ancestral reconstruction is likely to yield a structure that remains similar to those of its descendants over short evolutionary timescales. To test this hypothesis, we analyzed 39 protein families from the OrthoMaM database (version 12; Allio et al., 2024), each containing an average of 172 extant species with protein lengths of ~300 amino acids. To support ASR with such a high number of species, a hierarchy-based approach was developed (see Methods). For each protein family, we inferred the ancestral root sequence using BetaReconstruct and compared its predicted structure with the structure predicted based on the ancestral sequences inferred using FastML and ARPIP. To assess structural similarity, we randomly sampled one representative species from each major clade (defined by taxonomic orders; usually 25 randomly selected proteins) and predicted their structures using the AlphaFold 3 webserver (Abramson et al., 2024). We then compared the predicted structure of each reconstructed ancestral sequence to the structures of its sampled descendants using TM-align (version 20190822; Y. Zhang & Skolnick, 2005) and Foldseek (van Kempen et al., 2024), which reports similarity scores including bit score, alignment length, and percent sequence identity. We additionally compared the structure of a randomly selected leaf to those of a set of randomly chosen leaves ("random leaf").

##### TM-align similarity

TM-align (Y. Zhang & Skolnick, 2005) performs a sequence-independent alignment of protein tertiary structures. Unlike methods that rely on local distance metrics, TM-align employs a heuristic algorithm to identify the optimal global superposition by maximizing the Template Modeling score (TM-score). This metric is normalized by the length of the target protein ("ttmscore"), or the query protein ("qtmscore"), making it insensitive to protein size and providing a more robust measure of global fold similarity than Root Mean Square Deviation ("RMSD"). The resulting similarity score ranges from 0 to 1, where a score above 0.5 generally indicates that the proteins share the same fold, effectively capturing the topological relationship between the predicted models and the target orthologs.

##### Foldseek similarity

Foldseek (van Kempen et al., 2024) calculates structural similarity by transforming the three-dimensional structures into sequences over a structural alphabet, designed to encode tertiary interactions, and computing a distance between the resulting structure-derived sequences. The similarity score reflects both geometric alignment and sequence conservation within the structural alphabet. This approach allows Foldseek to efficiently detect structural similarity between proteins.

Table S5 summarizes the results across 39 protein families, reporting the mean structural similarities between the predicted ancestral structures and those of randomly selected leaves. BetaReconstruct achieves competitive scores compared to FastML and ARPIP. To assess the statistical significance of the differences between BetaReconstruct and either FastML or ARPIP, we conducted paired t-tests, which showed that none of the differences were statistically significant except for a slightly higher score of BetaReconstruct compared to ARPIP when comparing the fraction of identical matches (fident; paired t-test; $p < 0.03$; Table S5b). The structural similarity between the ancestors and a sample of leaves was higher than the structural similarity between two leaves ("Random leaf"). Overall, these results suggest that the structures predicted by BetaReconstruct are biologically plausible.

*Table S5: Comparing the structures of inferred ancestral sequences. The ancestral sequences were predicted using BetaReconstruct, FastML, or ARPIP. The predicted ancestral structures were compared to a random set of leaf structures. "Random leaf" represents the structural comparison between two leaves. Structural differences were evaluated using TM-align (a) and Foldseek (b). TM-align parameters: alnlen (number of aligned columns), RMSD (Root Mean Square Deviation), qtmscore (TM-score normalized by the query length), ttmscore (TM-score normalized by the target length), and coverage (proportion of the protein sequence successfully superimposed and aligned in 3D space). Foldseek parameters: fident (fraction of identical matches; measuring amino acid identity), alnlen, evalue (E-value), bits (bit score), alntmscore (TM-score of the alignment; Xu & Zhang, 2010), qtmscore, ttmscore, lddt (average LDDT of the alignment; Mariani et al., 2013), and prob (estimated probability for query and target to be homologous). Higher scores indicate better performance for all columns, except for the evalue, where lower values indicate better matches.*

*(a)*

| | alnlen | RMSD | qtmscore | ttmscore | Coverage |
|---|---|---|---|---|---|
| **BetaReconstruct** | 280.025 | 1.676 | 0.852 | 0.858 | 0.904 |
| **ARPIP** | 288.264 | 1.659 | 0.855 | 0.86 | 0.907 |
| **FastML** | 280.644 | 1.726 | 0.855 | 0.86 | 0.912 |
| **Random leaf** | 182.257 | 1.432 | 0.707 | 0.703 | 0.757 |

*(b)*

| | fident | alnlen | evalue | bits | alntmscore | qtmscore | ttmscore | lddt | prob |
|---|---|---|---|---|---|---|---|---|---|
| **BetaReconstruct** | 0.958 | 300.589 | 0 | 2121.465 | 0.854 | 0.844 | 0.85 | 0.952 | 1 |
| **FastML** | 0.96 | 299.752 | 0 | 2130.087 | 0.858 | 0.848 | 0.852 | 0.953 | 1 |
| **ARPIP** | 0.953 | 311.436 | 0 | 2179.249 | 0.859 | 0.847 | 0.852 | 0.949 | 1 |
| **Random leaf** | 0.941 | 298.649 | 0 | 2084.734 | 0.858 | 0.848 | 0.842 | 0.952 | 0.997 |

#### Supplementary Information S6: Training hyperparameters

For each of the three configurations (Configurations 1, 2, and 3), we trained two models using different learning rates: $10^{-4}$ and $10^{-5}$. In addition, the models were trained using mixed-precision, i.e., specific calculations were performed in 16-bit rather than 32-bit, which reduces the memory footprint and the training time (Micikevicius et al., 2018). The scheduler is “cosine”, the batch size is 128, and the maximum number of tokens is set to 2,048. We use a warmup period of 512 steps, and the maximum number of steps is 20,480. For the mammalian data, the maximum steps hyperparameter is 2,500, with 250 warmup steps.
